# Effects of Microplastics and Drought on Ecosystem Functions and Multifunctionality

**DOI:** 10.1101/2020.07.26.221929

**Authors:** Yudi M. Lozano, Carlos A. Aguilar-Trigueros, Gabriela Onandia, Stefanie Maaß, Tingting Zhao, Matthias C. Rillig

**Affiliations:** Freie Universität Berlin, Institute of Biology, Plant Ecology. D-14195 Berlin, Germany; Berlin-Brandenburg Institute of Advanced Biodiversity Research (BBIB), D-14195, Berlin, Germany; Leibniz Centre for Agricultural Landscape Research (ZALF), Dimensionality Assessment and Reduction, D-15374, Müncheberg, Germany; Universität Potsdam, Institute of Biochemistry and Biology, Plant Ecology and Nature Conservation, Am Mühlenberg 3, D-14476 Potsdam

**Keywords:** Enzymatic activities, global change ecology, grasslands ecosystem, litter decomposition, nutrient cycling, nutrient leaching, soil pH, soil aggregation, soil respiration

## Abstract

1. Microplastics in soils have become an important threat for terrestrial systems, which can be exacerbated by drought as microplastics may affect soil water content. Thus, the interaction between these two factors may alter ecosystem functions such as litter decomposition, stability of soil aggregates, as well as functions related to nutrient cycling. Despite this potential interaction, we know relatively little about how microplastics, under different soil water conditions, affect ecosystem functions and ecosystem multifunctionality.
2. To address this gap, we carried out a controlled-environment study using grassland plant communities. We applied the two factors microplastic fibers (absent, present) and soil water conditions (well-watered, drought), in all possible combinations in a factorial experiment. At harvest, we measured multiple ecosystem functions linked to nutrient cycling, litter decomposition, and soil aggregation and as terrestrial systems provide these functions simultaneously, we also assessed ecosystem multifunctionality.
3. Our results showed that the interaction between microplastic fibers and drought affected ecosystem functions and multifunctionality. Overall, drought had negatively affected nutrient cycling by decreasing potential enzymatic activities and increasing nutrient leaching, while microplastic fibers had a positive impact on soil aggregation and nutrient retention by diminishing nutrient leaching. Microplastic fibers also impacted enzymatic activities, soil respiration and ecosystem multifunctionality, but importantly, the direction of these effects depended on soil water status (i.e., they decreased under well watered conditions, but tended to increase or had similar effects under drought conditions). Litter decomposition had a contrary pattern.
4. *Synthesis and applications*. As soil water content is affected by climate change, our results suggest that areas with sufficiency of water would be negatively affected in their ecosystem functioning as microplastics increase in the soil; however, in areas subjected to drought, microplastics would have a neutral or slightly positive effect on ecosystem functioning.

## 1. INTRODUCTION

Microplastics are a group of polymer-based particles with a diameter under 5 mm (Hidalgo-Ruz et al., 2012), which occur in many shapes, and possess a high physical and chemical diversity (Helmberger et al., 2020, Rillig, Lehmann, & Ryo 2019). These particles can originate from many sources, including tire abrasion, the loss of fibers from synthetic textiles during washing, or the environmental degradation of larger plastic objects (Boucher & Friot, 2017). In addition, many plastics are already produced as microplastics (primary microplastics), e.g. for use in the cosmetics industry (Boucher & Friot, 2017). Therefore, microplastics are ubiquitous around the globe and may pollute not only oceans but also terrestrial systems through soil amendments, plastic mulching, irrigation, flooding, atmospheric input and littering or street runoff (Bläsing & Amelung, 2018; Rillig, 2012; de Souza Machado et al., 2018).

Our knowledge about microplastic effects on ecosystem functions is limited (Rillig and Lehmann, 2020) and potential interactive effects of microplastics with soil water availability are unknown. Among microplastics, microfibers are considered one of the most abundant microplastic types in the soil (Zhang and Liu, 2018, Dris et al., 2015), and these can potentially affect soil-water dynamics due to their linear shape, size and flexibility. For instance, microplastic fibers can enhance soil water holding capacity and so lead to the retention of water for longer periods (de Souza Machado et al., 2019), thus altering soil water conditions, and potentially influencing ecosystem functions. Indeed, microplastic fibers may promote plant growth and other processes (de Souza Machado et al., 2019), and this could alleviate drought conditions promoting plant productivity at the community level (Lozano and Rillig, 2020). All of this suggests that microplastic effects on ecosystem functionality may be exacerbated when other global change drivers, such as drought, come into play.

This potential interaction between microplastics in the soil and drought can affect multiple ecosystem functions involved in nutrient cycling, litter decomposition or soil aggregation. However, research on how microplastics and drought affect such functions has been limited. For example, nutrient cycling and energy flows are closely related to soil enzymes produced by microbes and plants (Stark et al., 2014), and enzymatic activity is highly influenced by environmental factors such as soil pH, nutrient availability and soil water content (Paul & Clark, 1989). By altering these factors, microplastics may potentially affect soil enzymatic activities. Indeed, there is evidence for microplastic influencing some enzymes: microplastics can stimulate or inhibit the activity of fluorescein diacetate hydrolase depending on the polymer type (de Souza Machado et al., 2019; Fei et al., 2020, Liu et al., 2017), or stimulate phenol oxidase (Liu et al., 2017), urease and acid phosphatase activities (Fei et al., 2020). In contrast, data on the effect that microplastic may have on key enzymes related to C, N, P-cycling (such as ß-glucosidase and ß-D-cellobiosidase involved in cellulose degradation, or ß-glucosaminidase involved in chitin degradation) are missing or limited (as in the case of phosphatases).

Litter decomposition is also a key ecosystem function with a crucial role in carbon cycling (Schmidt et al., 2011). This process depends on many factors including soil water content, litter quality and the decomposer community (Paul & Clark, 1989). Microplastics may directly affect decomposition by modifying some of these factors, or indirectly through its effects on soil aggregation (a function that is highly correlated with decomposition). So far, empirical evidence of the effect of microplastics on litter decomposition is sparse (Barreto et al., 2020), and we know even less about how decomposition might be affected under different water regimes (e.g., well-watered, drought conditions). Similarly, there are few data on microplastic impacts on soil aggregation, a key ecosystem function (Giling et al., 2019) which is also affected by biotic and abiotic factors (Bronick & Lal, 2005), and influences soil water dynamics and soil carbon storage (Peng et al., 2015). Microplastics may affect soil aggregation processes as they could reduce the stability of soil aggregates by affecting soil biota (Lehmann et al., 2019, Liang et al., 2019, de Souza Machado et al., 2019). Microplastics can also promote soil aggregation by helping to entangle soil particles (Rillig, Ryo et al., 2019) and by keeping the water in the soil for longer (de Souza Machado et al., 2019). This would counteract the negative effects that drought may have on soil aggregation (Zhang et al., 2018).

The trends summarized above not only illustrate the scarce knowledge about the effects of microplastic on terrestrial ecosystem functions, but also suggest the potential link between microplastics and drought as changes in soil water conditions may exacerbate the magnitude of microplastic effects and its direction (positive or negative), depending on the function measured. The net effect of each ecosystem function can alter the overall functioning of the soil. Given this heterogeneity of effects, and that ecosystem functioning is inherently multidimensional, addressing how microplastic influence multifunctionality (defined as the ability of an ecosystem to deliver multiple functions simultaneously (Hector & Bagchi, 2007)) could generate an integrative understanding of the terrestrial systems response to this global change driver.

To address these questions, we established microcosms, containing plant communities, on which we assessed the effect of microplastic fiber addition and drought in a factorial design given that we expect microplastic fibers to affect soil-water dynamics, on different ecosystem functions related to nutrient cycling, soil aggregation, decomposition, (Giling et al., 2019) and on ecosystem multifunctionality. We expected that microplastic fibers would affect single ecosystem functions and ecosystem multifunctionality in a positive or negative way depending on soil water conditions.

## 2. MATERIALS AND METHODS

### 2.1. Microplastics and soil preparation

In Dedelow, Brandenburg, Germany (53º 37’ N, 13º 77’ W), we collected dry sandy loam soil from grasslands communities (0.07% N, 0.77% C, pH 6.66). Soil was sieved (4 mm mesh size), homogenized and mixed with microplastic fibers at a concentration of 0.4%. This concentration aimed to simulate low to medium level of microplastic pollutions, since in soils of highly polluted areas a microplastic concentration up to ∼7% was observed (Fuller and Gautam, 2016). To do so, we manually cut with scissors polyester fibers (Rope Paraloc Mamutec polyester white, item number, 8442172, Hornbach.de) to generate microplastic fibers that had a length of 1.28 ± 0.03 mm. Twelve grams of microplastic fibers (∼763333 fibers g^-1^ microplastic) were mixed into 3 kg of soil for each pot. For each experimental unit, microplastic fibers were separated manually and mixed with the soil in a large container before placing into each individual pot, to help provide a homogeneous distribution of microplastic fibers throughout the soil and the intended microfiber concentration. Twenty experimental units (pots) were established. Half had soil with microplastic fibers, while the other half had soil without added microplastic fibers. Soil was mixed in all experimental units in order to provide the same level of disturbance. For additional details see Lozano and Rillig (2020).

### 2.2. Experimental setup

In May 2019 we established the experiment in a temperature-controlled glasshouse with a daylight period set at 12 h, 50 klx, a temperature regime at 22/18 ºC day/night, and a relative humidity of ∼40 %. We selected seven grassland plant species frequently co-occurring in Central Europe, which naturally grow in the same patch in dry grasslands in the Brandenburg region, Germany. Seeds of *Festuca brevipila, Holcus lanatus, Calamagrostis epigejos, Achillea millefolium, Hieracium pilosella, Plantago lanceolata* and *Potentilla argentea*, were obtained from a commercial supplier in the region (Rieger-Hofmann GmbH, Blaufelden, Germany) in order to shape a plant community typical of temperate grasslands ecosystems. Seeds were surface-sterilized with 10% sodium hypochlorite for 5 min and 75% ethanol for 2 minutes, thoroughly rinsed with sterile water and germinated in trays with sterile sand. Then, we randomly transplanted seedlings of similar size into pots (16 cm diameter, 16.5 cm height, 3L) where twenty-one holes were dug with a distance of 2.5 cm. This way, a plant community consisting of three individuals of each of the seven plant species was established in each pot. We will refer to plant species by their generic names from now on.

Pots were well-watered (100 ml twice a week) during the first three weeks of growth. Then, half of them were kept at ∼70% of soil water holding capacity (WHC) by adding 200 ml of water, while the other half were kept at ∼ 30% WHC by adding 50 ml of water. Pots were watered from the top twice a week for two months with distilled water. Previous assays showed that these amounts and frequency of watering keep the established WHC. We thus had 20 experimental units in a fully crossed orthogonal design that includes two microplastic fiber treatments (one with and the other without added microplastic fibers, also called “present” and “absent”) and two drought treatments (with and without drought, also called “drought” and “well-watered”), with five replicates each (n = 5). Pots were randomly distributed in the chamber and their position was shifted twice to homogenize environmental conditions experienced by each replicate during the experiment.

At harvest we measured eleven variables that capture aspects of decomposition, nutrient cycling and soil structure formation (litter decomposition, ß-glucosidase, ß-glucosaminidase, ß-D-cellobiosidase, phosphatase, soil respiration, water stable aggregates, leaching of NO_3_^-^, SO_4_^2-^, PO_4_^3-^, and soil pH; functions hereafter).

### 2.3. Measurement of soil ecosystem functions

#### Soil nutrient cyclin

In fresh soil, we measured four functions related to C, N and P cycling: activity of ß-glucosidase and ß-D-cellobiosidase (cellulose degradation), N-acetyl-ß-glucosaminidase (chitin degradation) hereafter ß-glucosaminidase, and phosphatase (organic phosphorus mineralization). Extracellular potential soil enzyme activities were measured from 1.0 g of soil by fluorometry as described in Bell et al. (2013).

#### Soil respiration

We took a 25 g soil subsample from each pot to measure soil respiration via an infrared gas analyzer. To do this, we placed the subsamples in individual 50 ml falcon tubes with modified lids that allow control of gas exchange via a rubber septum. We measured CO_2_ concentration (ppm) at two time points from these falcon tubes as described in Rillig, Ryo et al., 2019. The first time point was obtained after we flushed the tubes with CO_2_ free air for five minutes thus reflecting CO_2_ concentration at time 0. The second point was obtained after letting the tubes with the soil samples incubate at 25°C for 65 h. At both time points, we took a 1-mL air sample and injected it to an infrared gas analyzer (LiCOR-6400XT). We report soil respiration as the net CO_2_ production (in ppm) after the incubation period by subtracting the measurement from the first time point from that of the second.

#### Litter decomposition

We collected plant material from dry grasslands where our species naturally grow (see Onandia et al., 2019 for methodological details) and obtained a composite sample that reflected the proportion of plant biomass of each plant species in the field. Plant material was oven-dried at 60 ºC for 72 h, milled, and 0.75 mg were placed in 6×6 cm polyethylene terephthalate (PET, Sefar PET 1500, Farben-Frikell Berlin GmbH, Germany) bags with a mesh size of 49 µm. One litter bag was buried in each pot at 8 cm depth prior to seedling transplanting, and retrieved at harvest. Litter bags were stored at 4°C and processed within 2 weeks. Soil attached to the bags was carefully washed away using tap water and then, litter decomposition was estimated as mass loss after each bag was oven-dried at 60°C for 72 h.

#### Soil aggregatio

Water stable soil aggregates are a proxy measure of soil aggregation and were measured following a modified version of the method of Kemper and Rosenau (1986), as described in Lehmann et al., 2019. Briefly, 4.0 g of dried soil (<4 mm sieve) was placed on small sieves with a mesh size of 250 μm. Soil was rewetted with deionized water by capillarity and inserted into a sieving machine (Agrisearch Equipment, Eijkelkamp, Giesbeek, Netherlands) for 3 min. Agitation and re-wetting causes the treated aggregates to slake. We collected the soil left on the sieve (coarse matter + water stable fractions, also called dry matter) and then separated the coarse matter by crushing the aggregates and pushing the soil through the sieve. Dry matter and then coarse matter were dried at 60 °C for 24 h. Soil aggregation (i.e., water stable aggregates) was calculated as: WSA (%) = (Dry matter-coarse matter)/(4.0 g - coarse matter).

#### Soil nutrient leaching and pH

At harvest, pots were watered to saturate the soil to roughly 10% beyond the water holding capacity, simulating a rain event, to induce leaching. Leachate percolating through the soil column was collected from small outlets at the bottom of the pot and was assessed for nutrient concentrations (NO_3_^-^, SO4 ^2-^, PO4 ^3-^) using ion chromatography (Dionex ICS-1100, AS9-HC, Thermo Scientific Massachusetts, USA). Air-dried soils were extracted in deionized water for 1 h to achieve a 1:5 (v:v) soil: water solution and soil pH was determined with a Hanna pH-meter (Hanna Instruments GmbH, Vöhringen, Deutschland).

### 2.4. Assessing ecosystem multifunctionality

To calculate ecosystem multifunctionality we followed the ecosystem function multifunctionality method proposed by Manning et al. (2018). Briefly, we identified the clusters of 12 ecosystem functions (Figure S1), which included the soil functions measured in this study and total shoot mass (raw data obtained from Lozano and Rillig (2020)). This cluster analysis allowed us to give more even weights to the ecosystem functions as they are interrelated and shared drivers. We determined the number of clusters by the Elbow method, (Kassambara & Mundt, 2017) and weighted each of them equally, irrespective of the number of functions within each cluster. Four clusters were determined. Then, we calculated the standardized maximum for each function and placed the function data on a standardized scale. Thus, we standardized by the average of the top 10% values within the data and calculated ecosystem multifunctionality for each experimental unit using the threshold approach, in which each ecosystem function that exceeds 70 % of the standardized maximum contributed one to the ecosystem multifunctionality score. Additional calculations of ecosystem multifunctionality were done using a threshold of 30% and 50% (Figure S2, Table S1).

### 2.5. Statistical analyses

The experimental design was a fully crossed orthogonal design where microplastic fibers, drought, and the interaction were considered fixed factors. Each function was analyzed using linear models. Model residuals were checked to validate normality and variance homogeneity assumptions. We implemented the “varIdent” function to account for heterogeneity in the microplastic fiber treatment for ß-D-cellobiosidase, soil aggregation, and in the water treatment for soil respiration. The effect of microplastics and drought on the ecosystem multifunctionality index was analyzed using generalized linear models with a quasibinomial distribution and a logit link function to avoid overdispersion. We also assessed the contribution of each function to multifunctionality by using the down-weighting data after clustering and the metric “pmvd” from the package “relaimpo” (GrLmping, 2006). This metric is based on sequential R^2^s, but takes care of the dependence on orderings by weighted averages with data-dependent weights and also guarantees that a regressor with 0 estimated coefficient is assigned a relative importance of 0 (GrLmping, 2006). Statistical analyses were done with R version 3.5.3 (R Core Team, 2019). Results shown throughout the text are mean values ± 1 standard error (SE).

## 3. RESULTS

Ecosystem functions were affected by microplastic fibers, drought and their interaction (Table 1). While enzymatic activities and soil respiration were on average higher under well-watered than under drought conditions, these trends changed in the presence of microplastics, decreasing under well-watered conditions but increasing under drought. As for enzymatic activity, ß-glucosaminidase decreased by 29% with drought and was not affected by microplastic fibers (Table 1, Figure 1). ß-D-cellobiosidase decreased by 62% with drought (p = 0.02), while soil respiration was marginally affected by microplastic fibers and drought (p = 0.1). Phosphatase and ß-glucosidase were affected by the interaction between microplastic fibers and drought (p = 0.03, p = 0.1, respectively). Both decreased with microplastic fibers in soil by 27% and 17% under well-watered while increasing by 75% and 40% under drought conditions, respectively (Table 1, Figure 1). By contrast, litter decomposition increased with microplastic fibers by 6.4 % under well-watered conditions while decreasing by 6.6% under drought conditions (p = 0.09, Figure 1). Likewise, soil aggregation increased with microplastic fibers under both well-watered and drought conditions by 15 % and 21.7 %, respectively (p = 0.07). Overall, soil leachate nutrients increased with drought and decreased with microplastic fibers in the soil. Specifically, leachate NO_3_^-^ decreased by 70% with microplastic fibers under drought conditions (p = 0.01, Figure 1), a similar trend was found under watered conditions. Leachate SO4 ^2-^ decreased with microplastic fibers under either well-watered or drought conditions by 52% and 37%, respectively (p = 0.01). PO4 ^3-^ in leachate was not clearly affected by drought or microplastic fibers, while soil pH increased both with drought and microplastic fibers in the soil (p < 0.01, Figure 1).

**TABLE 1.** Results from linear models on eleven ecosystems functions and multifunctionality response to microplastic fibers (M), drought (D) and their interaction (M x D). Multifunctionality also included shoot mass (data extracted from Lozano and Rillig, 2020). Degrees of freedom of each factor (df =1). F values and p-values (in parentheses) are shown; p values <0.1 in bold. n = 5.

| Ecosystem functions | Microplastic<br>fibers (M) | Drought (D) | M x D |
| --- | --- | --- | --- |
| β-glucosaminidase | 0.14 (0.70) | 2.98 (0.10) | 1.08 (0.31) |
| β-glucosidase | 0.02 (0.89) | **6.88 (0.01)** | 2.31 (0.14) |
| Phosphatase | 0.07(0.79) | **3.55(0.07)** | **5.53 (0.03)** |
| β-D-cellobiosidase | 2.14 (0.16) | **6.32 (0.02)** | 1.49 (0.23) |
| Soil respiration | 2.49 (0.13) | 2.29 (0.14) | 1.37 (0.25) |
| Litter decomposition | 0.002 (0.95) | 0.88 (0.36) | **3.13 (0.09)** |
| Soil aggregation | **3.54 (0.07)** | 2.51(0.13) | 0.03(0.84) |
| $\text{NO}_3^-$ | **10.66 (0.004)** | **24.93 (0.0001)** | **7.85 (0.01)** |
| $\text{PO}_4^{3-}$ | 0.36 (0.55) | 0.25 (0.62) | 0.08 (0.77) |
| $\text{SO}_4^{2-}$ | **6.75 (0.01)** | **3.66 (0.07)** | 0.00 (0.99) |
| pH | **12.38 (0.002)** | **9.14 (0.008)** | 0.47 (0.50) |
| Multifunctionality | 3.16 (0.09) | 3.02 (0.10) | **7.23 (0.01)** |

**FIGURE 1.**
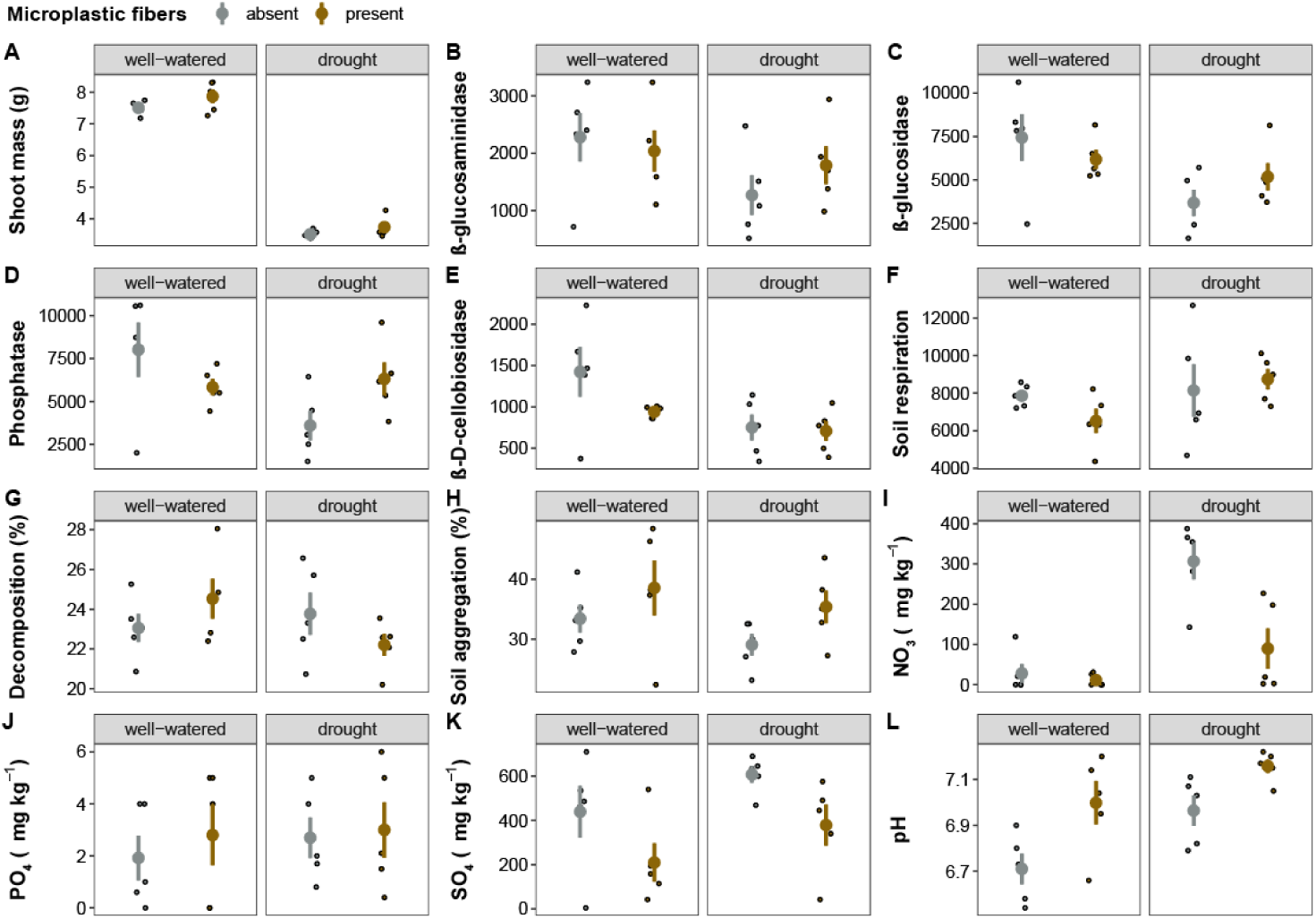
Microplastic fibers and drought effects on twelve ecosystem functions. Mean and standard error are represented. Data points are shown as circles. Enzymes and soil respiration units (µmol g^-1^ dry soil hr^-1^, ppm). P-values in Table 1; n = 5.

Ecosystem multifunctionality was affected by the interaction between microplastic fibers and drought (Table 1, Figure 2). That is, the effect of microplastics on ecosystem multifunctionality strongly depended on the drought treatment (p = 0.01): under well-watered conditions, microplastic fibers addition to the soil decreased multifunctionality, while under drought conditions, microplastic addition did not affect multifunctionality (Figure 2).

**FIGURE 2.**
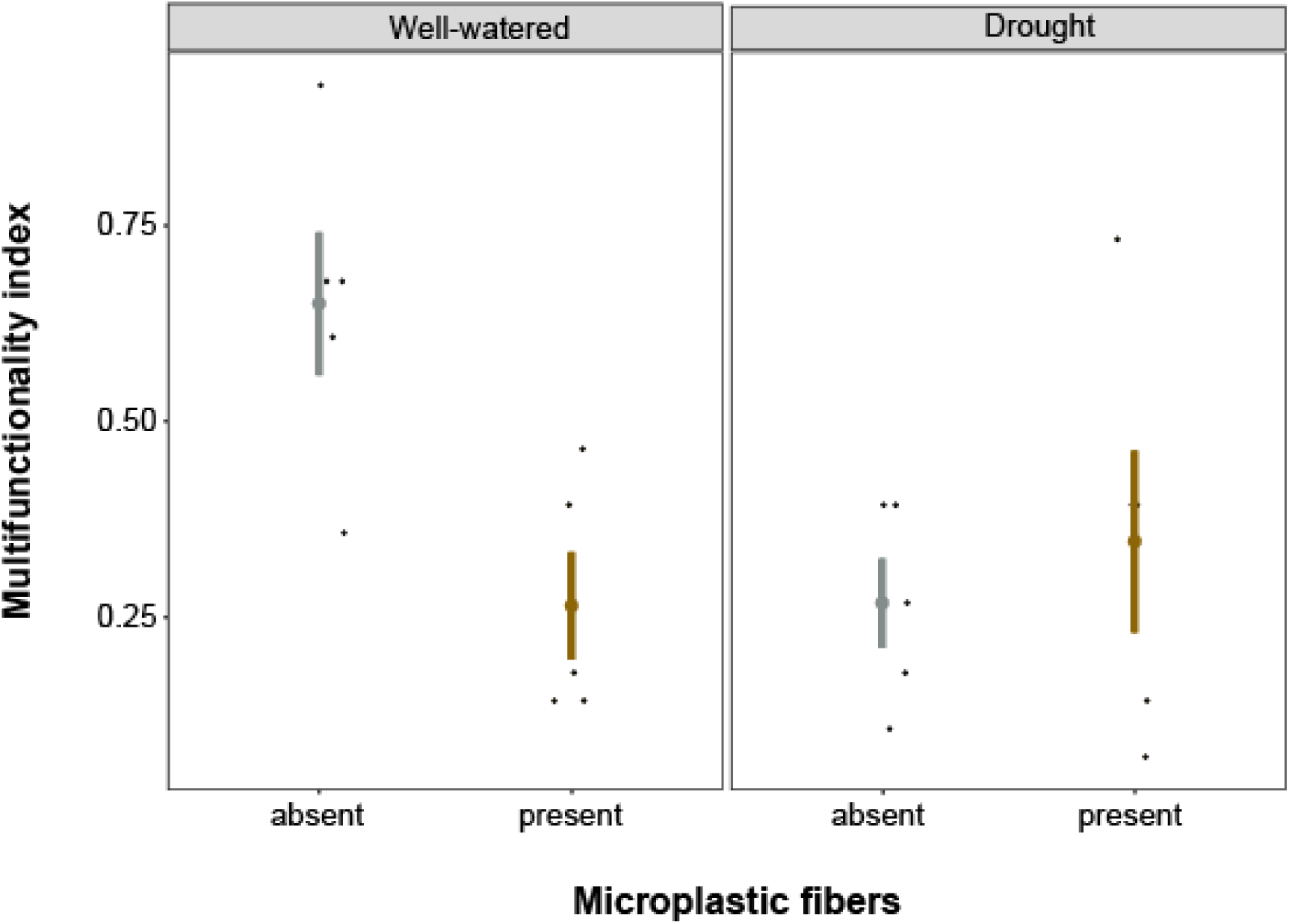
Microplastic fibers and drought effects on ecosystem multifunctionality. Mean and standard error are represented. Data points are shown as circles; P-values in Table 1; n = 5.

Different thresholds when calculating multifunctionality showed similar trends (Figure S2, see Table S1 for statistical results). The analysis of the relative importance of each function showed that ß-glucosidase (31.87 %), soil respiration (25.65 %), phosphatase (11.14 %), pH (9.16 %), SO_4_^2-^ (8.84 %), ß-D-cellobiosidase (3.03 %), ß-glucosaminidase (2.88 %), shoot mass (1.88 %), PO_4_^3-^ (1.67 %), soil aggregation (1.63 %), litter decomposition (1.56 %), NO_3_^-^ (0.62%) contributed in this order to multifunctionality (R^2^ = 91.53 %, Figure 3).

**FIGURE 3.**
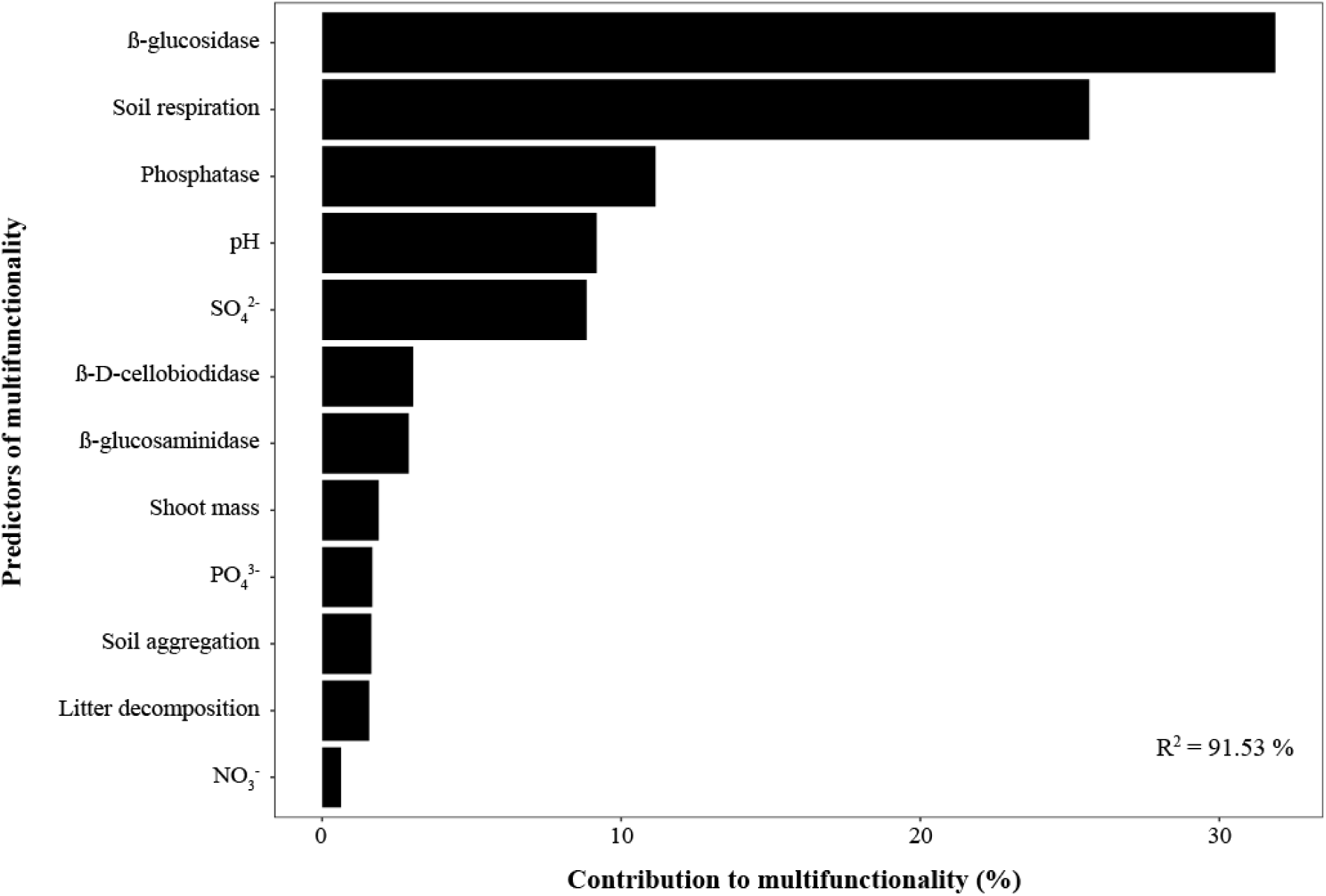
Relative importance of each predictor to multifunctionality. The proportionate contribution of each function considered both its direct effect (i.e., its correlation with multifunctionality) and its effect when combined with the other variables in the regression equation. The metrics “pmvd” was used for the calculation and the down-weighting via the cluster was taken into account.

## 4. DISCUSSION

As hypothesized, microplastic fibers and drought affected ecosystem functions linked with soil aggregation, nutrient cycling and decomposition as well as ecosystem multifunctionality. Overall, drought had a negative impact on ecosystem functions, while the impact of microplastic fibers depended on the soil water status and the function considered. Below, we discuss likely mechanisms behind these complex outcomes.

### 4.1. Soil aggregation increased with microplastic fibers irrespective of drought

Microplastic fibers promoted soil aggregation either under well-watered or drought conditions, likely due to positive effects of fibers on soil bulk density, aeration and water retention (de Souza Machado et al., 2019), which may promote root growth (Lozano & Rillig, 2020) and hyphal extension (Elliot & Coleman, 1988; Wang et al., 2017). Therefore, roots, hyphae and microplastic fibers might together have helped entangle soil particles, thus promoting soil aggregation. In addition, microbial communities might have shifted, and this may also have contributed to the observed soil aggregation response.

### 4.2. Microplastic fibers reduce soil enzyme activity and soil respiration only under well watered conditions

We observed that microplastic fibers affected potential enzymatic activities and soil respiration depending on soil water conditions. That is, under drought, enzymes and soil respiration increased when microplastic fibers were added, probably because soil water content and aeration may increase with microplastic fibers (de Souza Machado et al., 2019; Rillig et al., 2019), which in turn may promote microbial activity (Nannipieri et al., 2002, Alster et al., 2013, Sanaullah et al., 2011). By contrast, under well-watered conditions, enzymes and soil respiration decreased with microfibers in the soil, probably linked with a decline in soil microbial community richness and diversity as seen by Fei et al. (2020), a negative effect that could be exacerbated if microfibers may release harmful contaminants into the soil (Rillig, 2012; Wang et al., 2019).

### 4.3. Microplastic fibers increase litter decomposition under well-watered conditions

Litter decomposition increased under well-watered conditions when microplastic fibers were added. Our results suggest that the increase in litter decomposition may be related to an increase in soil aggregation. Soil aggregation promotes oxygen diffusion within larger soil pores and regulates water flow, which in turn stimulate microbial activity (Six et al., 2004) promoting litter decomposition. In addition, soil pH, a parameter influenced by soil aggregation (Jiang et al., 2013), that affects soil microbial community structure (Fierer & Jackson, 2006), could also have played a role. In fact, recent research found that an increase in litter decomposition was linked with better soil aggregation (Yang et al., 2019). Our results suggest that microplastics, through effects on litter decomposition may have large consequences for ecosystem C stocks and fluxes, as changes in litter decomposition may influence the feedback to the atmosphere from terrestrial ecosystems.

### 4.4. Microplastics fibers reduce soil nutrient leaching

Nutrient leaching, after a simulated rain event, increased under drought but decreased when microplastic fibers were added to the soil. Drought conditions might have led to the formation of cracks as preferential flow paths in the soil, increasing the leaching of nutrients when the soils were rewetted. In support of this, in fertilized soils the leachate NO3 ^-^ was threefold higher under drought than under non-drought conditions (Klaus et al., 2020). Nutrient leaching is also known to be related to change in the structure of plant and microbial communities (Mueller et al., 2013), biotic factors that are indeed affected by drought (Lozano et al., 2019, Fitzpatrick et al., 2018). Likewise, we observed that leachate PO_4_^3-^ was not affected by drought, most likely because phosphates are more strongly bound to soil particles than nitrate or sulphate (Paul & Clark, 1989). By contrast, nutrient leaching decreased with microplastic fibers (i.e., more nutrient retention). This can be related to the positive effect that microfibers had on soil aggregation, which may have increased the soil capacity to retain nutrients. This positive relation between soil nutrients retention and soil aggregation has been reported by Liu, Han, & Zhang (2019).

### 4.5. Microplastic fibers and drought effects on ecosystem multifunctionality and ecosystem services

Our results showed that microplastic fibers and drought impacted not only single functions but also multifunctionality, and that such impact depended on the interaction between these two global change factors. Specifically, with the addition of microplastic fibers, ecosystem multifunctionality decreased under well-watered conditions, while giving rise to similar functioning under drought conditions. This trend mirrors the one observed for nutrient cycling functions (i.e., ß-glucosidase, soil respiration), as they are the ones that contribute most to multifunctionality. Thus, this result highlights the importance of considering nutrient cycling functions when managing microplastics in soils.

Our results showed that two global change drivers (i.e., microplastics and drought) influence ecosystem functions and multifunctionality, which in turn may affect ecosystem services (Manning et al., 2018; Díaz et al., 2018) and thus impact various aspects of human well-being. In the short term, microplastic fibers may contribute to plant productivity or soil aggregation; however, we do not currently know what the long-term responses will be, as additional factors could come into play. Indeed, microplastic fibers may release harmful chemical substances into the soil (Fred-Ahmadu et al., 2020) and affect nutrient cycling processes, with consequences for soil quality, and thus on the provision of different services, such as food and water (MEA, 2005). This becomes relevant as agricultural lands are often managed with sewage sludge or compost, which contains a large amount of microplastic fibers (Wang et al., 2019; Weithmann et al., 2018).

As microplastics may come into the soil in different shapes (Rillig et al., 2019) and polymer types (Helmberger et al., 2020), it is important to understand how different microplastic types may affect ecosystem functionality. However, our findings provide clear empirical evidence that microplastics in soil affect ecosystem multifunctionality of terrestrial ecosystems, a phenomenon that may be strongly affected in future scenarios of global change, as changes in water regime are projected to occur in many areas worldwide. Our results also highlight the potential of microplastic to affect Earth system feedbacks of terrestrial ecosystems, especially via observed changes in litter decomposition, respiration fluxes and soil aggregation.

## ACKNOWLEDGMENTS

The work was funded by the German Federal Ministry of Education and research (BMBF) within the collaborative Project “Bridging in Biodiversity Science (BIBS-phase 2)” (funding number 01LC1501A). MCR additionally acknowledges funding through an ERC Advanced Grant (694368). We thank Tydings Morrison, Carlos Acame and Jenny M. Ospina for their assistance in the experimental setup and data collection. We thank Walter Waldman for helpful discussions about microplastics properties, Arthur Gessler for valuable advice about decomposition assessment and Dr. Peter Manning for helpful comments on earlier versions of this manuscript. The authors declare no competing financial interest.

## DATA AVAILABILITY STATEMENT

We will not be archiving data because all data used are present in the manuscript.

## AUTHOR CONTRIBUTIONS

YML, CAAT, GO and MCR conceived the ideas and designed methodology; YML, CAAT, GO and SM established and maintained the experiment in the greenhouse; ZTT analyzed the soil enzymatic activities. YML analyzed the data and wrote the first draft of this manuscript. All authors contributed to the final version and gave final approval for publication.

