## Supplemental files for "Effects of Microplastics and Drought on Ecosystem Functions and Multifunctionality"

**SUPPORTING INFORMATION**

**Figure S1.** Cluster analysis of twelve functions used to measure ecosystem multifunctionality

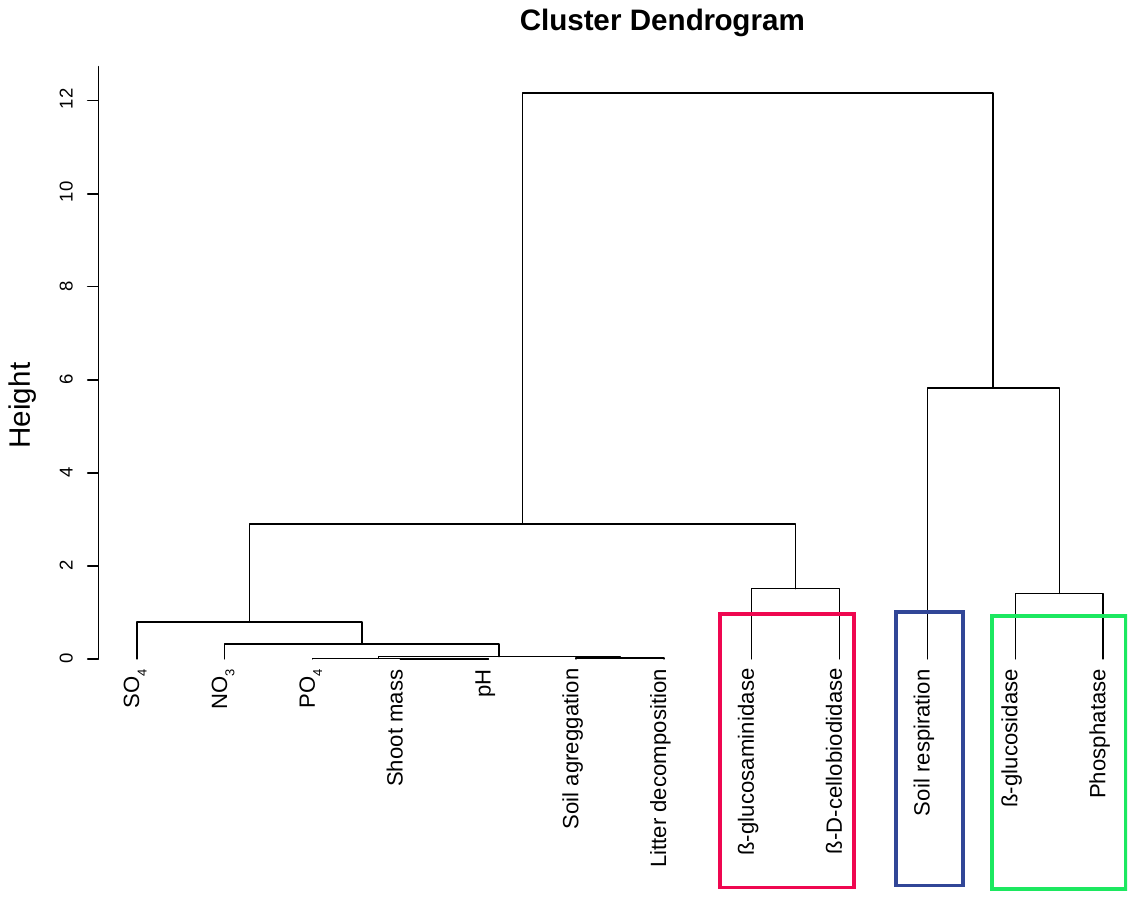

**Figure S2.** Microplastic fibers and drought effects on ecosystem multifunctionality. Mean and standard error are represented. Threshold at 30 and 50%. See statistical results in Table S1. Data points are shown as circles; n = 5.

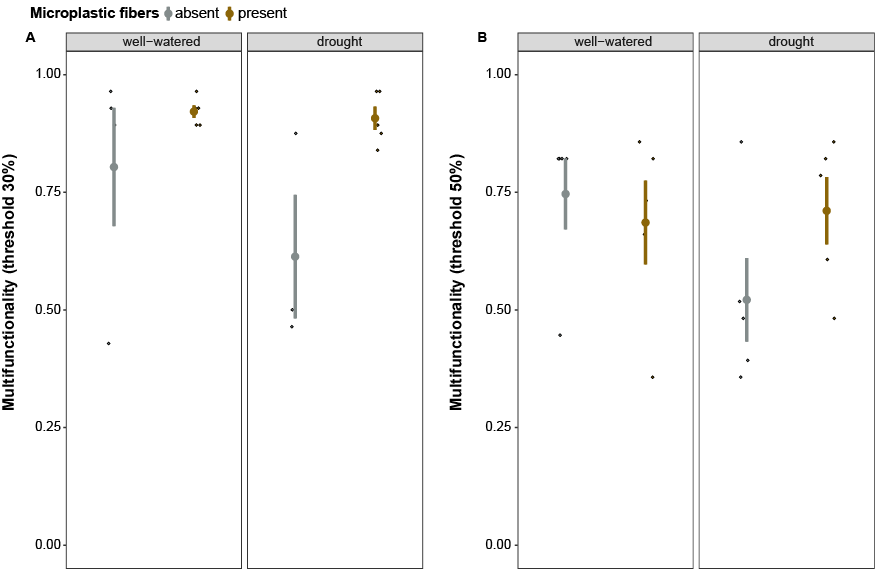

Table S1. Results from generalized linear models on ecosystem multifunctionality response to microplastic fibers (M), drought (D) and their interaction (M x D) at a threshold of 30% at 50%. n = 5.

|  |  | Threshold 30% | | Threshold 50% | |
| --- | --- | --- | --- | --- | --- |
|  | df | F value | P value | F value | P value |
| Microplastic fibers (M) | 1 | 3.28 | 0.08 | 0.62 | 0.44 |
| Drought (D) | 1 | 0.004 | 0.94 | 1.50 | 0.23 |
| M x D | 1 | 0.34 | 0.56 | 2.35 | 0.14 |
